# TNIK maintains a MYC-driven partial EMT state that supports proliferation and evasion of senescence in lung squamous cell carcinoma

**DOI:** 10.64898/2026.08.28.747625

**Authors:** Mohaddase Hamidi, Kenneth O Omolo, Korrey W Hart, Shrey Sitaram, Yan Zhou, Pedro Torres-Ayuso

## Abstract

Lung squamous cell carcinoma (LUSC) is an aggressive malignancy characterized by high cellular plasticity and few targeted treatment options. TNIK overexpression is common in LUSC and promotes tumor growth, with TNIK inhibition sensitizing LUSC to radiotherapy, though the underlying mechanisms are not well defined. Through transcriptomic analyses and functional assays, we identified TNIK as a regulator of a MYC-dependent transcriptional network that coordinates epithelial-mesenchymal plasticity and cell proliferation in LUSC. Depletion of TNIK reprogrammed LUSC cells from a hybrid epithelial/mesenchymal state towards an epithelial, senescent-like state characterized by reduced cell migration, invasion, reduced DNA synthesis, and enhanced β-galactosidase activity. Using a small-molecule screen approach, we found that TNIK inhibitors cooperated with agents suppressing the histone methyltransferase and MYC binding partner EZH2, which further suppressed partial epithelial-to-mesenchymal transition (pEMT). Mechanistically, we identified MYC as a key downstream TNIK effector in LUSC cells: MYC depletion phenocopied the effects of TNIK loss on pEMT and senescence, and restoring MYC expression bypassed the effects of TNIK depletion. Collectively, these results implicate TNIK in the mechanisms linking epithelial-mesenchymal plasticity with proliferation and evasion of senescence and provide insights into future strategies for the clinical deployment of TNIK inhibitors in LUSC and other TNIK-dependent malignancies.

## INTRODUCTION

Despite major efforts in therapeutic development, lung cancer remains the leading cause of cancer-related mortality worldwide and continues to be associated with poor clinical outcomes [1]. Lung squamous cell carcinoma (LUSC), the second most prevalent type of lung cancer worldwide, is characterized by high rates of relapse and limited therapeutic opportunities [2–4]. Unlike lung adenocarcinoma, LUSC generally lacks targetable driver mutations, although strategies targeting mTORC1/2 or BET proteins are under investigation [2, 5, 6]. Instead, LUSC is typically treated with combinations of chemoradiotherapy and immune checkpoint blockade, which cause damaging side effects; further, many patients have intrinsic resistance to treatment [3]. These facts highlight the need for improved understanding of the mechanisms sustaining LUSC progression and therapeutic resistance [7].

We recently identified the serine/threonine protein kinase TNIK (TRAF2- and NCK-interacting kinase) as a therapeutic vulnerability in a subset of LUSC [8]. TNIK is commonly upregulated by copy-number increase (as part of an amplicon on chromosome 3q) or by increased gene transcription in several tumor types [8–13]. Our work in LUSC defined the tumor suppressor MERLIN/NF2 as a critical TNIK substrate, and showed that TNIK-dependent phosphorylation of MERLIN led to activation of downstream effectors FAK and YAP, which was important for sustaining LUSC tumor growth [8]. Indicating a broader oncogenic activity of TNIK, studies in colorectal and ovarian cancers have defined additional TNIK-regulated pathways, including the WNT/β-catenin pathway [10, 13–15]. Importantly, we and others have found that small-molecule inhibitors of TNIK, such as NCB-0846, control the growth of tumors expressing high levels of TNIK and sensitize LUSC to radiotherapy [8, 10, 12, 13, 16]. More fully defining the mechanisms by which TNIK contributes to therapeutic resistance would be valuable for identifying ways to clinically leverage TNIK inhibitors to improve patient care.

Phenotypic plasticity, particularly manifested as epithelial-to-mesenchymal transition (EMT), is a major contributor to therapy resistance in LUSC. While epithelial or mesenchymal identities were initially considered as binary alternatives, cancer cells are now known to commonly undergo partial EMT (pEMT) and reside in intermediate or hybrid epithelial (E)/mesenchymal (M) states, which are associated with enhanced metastatic potential, stemness, and treatment resistance [17–21]. pEMT is promoted and sustained by the coordinated activity of multiple signaling pathways, including WNT/β-catenin, TGF-β, YAP/TAZ, NOTCH, and MAPK [19, 22], which converge on EMT transcription factors such as SNAIL (*SNAI1)*, SLUG (*SNAI2)*, TWIST1/2, and ZEB1/2. However, the interactions between signaling pathways and transcriptional networks that drive pEMT remain incompletely defined, thereby hindering the development of effective strategies to block or reverse pEMT and reprogram hybrid E/M cell states toward a therapy-responsive state [21, 23].

In this study, using RNA-seq of TNIK-depleted LUSC cells, we identified TNIK as an upstream regulator of multiple pEMT-driving transcriptional programs. Pathway analysis nominated MYC as a convergent target of multiple TNIK-regulated pathways. Using NCB-0846, supported by genetic manipulation of TNIK and MYC, we have confirmed that this core signaling axis regulates hybrid E/M cell states, phenotypic plasticity, and treatment resistance in LUSC.

## MATERIAL AND METHODS

### Cell lines

NCI-H520 (RRID: CVCL_1566) and SW900 cells (RRID: CVCL_1731) were obtained from ATCC. NCI-H520 and SW900 cells with doxycycline-inducible expression of pLKO-Tet-On shRNA control and TNIK-targeting constructs have been previously described [8]. Cell culture reagents and conditions are summarized in Supplementary Methods.

### RT-qPCR

Power SYBR Green RNA-to-CT 1-Step Kit (Applied Biosystems; Thermo Scientific, cat. 4389986) was used for RT-qPCR. The primers used are listed in **Supplementary Table 1**. Samples were run on a QuantStudio^TM^ 3 Real-Time PCR system (Applied Biosystems, RRID: SCR_018712) with the following program: reverse transcription (30 minutes, 48°C), activation of DNA polymerase (10 minutes, 95°C), PCR amplification (40 cycles of denaturing at 95°C for 15 seconds and annealing/extension at 60°C for one minute). SDS software was used to compute CT values. Relative gene expression was analyzed using the ΔΔCt method [24].

### Flow Cytometry Analyses

Fluorescence-activated cell sorting was used to evaluate cell proliferation, cell cycle phase distribution, and (E-Cadherin, EpCAM, vimentin, CD44) expression. Samples were analyzed in a BD FACSymphony A5 SE analyzer and data processed in FlowJo (v.10 or higher). Additional details are provided in Supplementary Methods.

### Statistical analysis

All experiments were performed in at least three independent biological replicates. Data are expressed as mean ± standard deviation (SD). Statistical significance was determined using Student’s t-test or one-way ANOVA as indicated in the corresponding figure legends. Statistical analyses were performed in Prism 11 (GraphPad Software, Boston, MA, RRID: SCR_002798). Significance was set at p = 0.05.

## RESULTS

### TNIK loss drives epithelial reprogramming and attenuates cell migration and invasion in LUSC

To identify the mechanisms by which TNIK contributes to LUSC tumor progression, we conducted RNA sequencing (RNA-seq) analyses on a panel of four LUSC cell lines (SW900, NCI-H520, LK-2, and Lc-1-sq) characterized by high TNIK protein levels and sensitivity to TNIK knockdown or inhibition [8]. HCC15 cells, which show no detectable TNIK expression and are not affected by TNIK depletion or inhibition [8], were used as a control. Ingenuity Pathway Analysis (IPA) of differentially expressed genes in LUSC cells after 72 hours of TNIK depletion versus corresponding control shRNA-expressing cells revealed a significant downregulation of transcriptional programs associated with cell migration, invasion, and pluripotency (**Fig. 1A**). Concurrently, transcriptional programs associated with cellular senescence and cell death were significantly upregulated in TNIK-depleted LUSC cells (**Fig. 1A**). The transcriptional changes observed in TNIK-depleted cells were consistent with cell state reprogramming toward an E, less stem-like state and led us to hypothesize that TNIK might support pEMT in LUSC cells.

**Figure 1.**
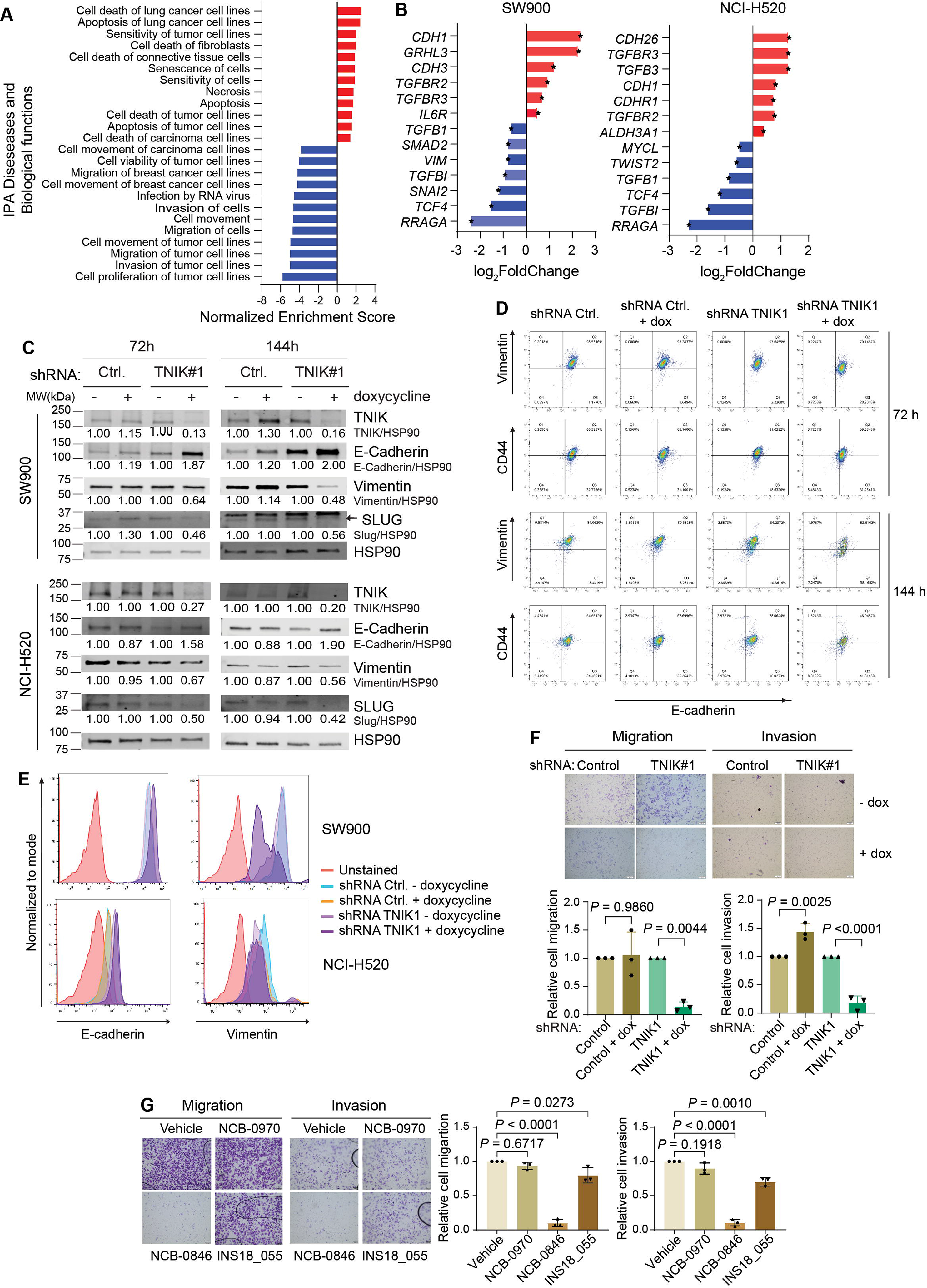
TNIK loss drives epithelial reprogramming and attenuates cell migration and invasion in LUSC cells. **A**, Ingenuity Pathway Analysis (IPA) analysis of RNA-seq from TNIK-depleted LUSC cells (SW900, NCI-H520, LK-2, and LC1/SQ) shows that TNIK loss significantly (p-adj < 0.05) downregulates (blue) biological pathways linked to cell plasticity, specifically EMT, while activating (red) senescence- and cell death-associated programs. **B**, Relative expression (Log_2_-fold change) of representative genes contributing to the enrichment of pEMT-related signatures in SW900 and NCI-H520 cells. Red and blue bars indicate significantly (p-adj < 0.05) upregulated and downregulated genes, respectively. **C**, Western blot analysis of TNIK, E-cadherin, vimentin, and SLUG *(SNAI2)* in whole-cell extracts from TNIK-depleted (dox-induced *TNIK* shRNA, 1 μg/mL, 72 or 144 hours) SW900 (top) or NCI-H520 (bottom) cells. HSP90 was used as a loading control. Protein level (band intensity) relative to HSP90 were normalized to untreated (minus dox) samples. The blot is representative of *n* = 3 independent experiments. **D**, Fluorescence-activated cell sorting analysis of E-cadherin, vimentin, and CD44 in SW900 cells after TNIK knockdown (dox-induced *TNIK* shRNA, 1 μg/mL, 72 or 144 hours). Hybrid E/M cells are defined as E-cadherin: vimentin or E-cadherin: CD44 double-positive cells. The graphs are representative of *n* = 3 independent experiments. **E**, Flow cytometry analysis of mean fluorescence intensity (MFI) values of E-Cadherin and vimentin expression in SW900 (top) and NCI-H520 (bottom) following depletion of TNIK (dox-induced, 1 μg/mL, 72 hours). The histogram is representative of *n* = 3 independent experiments. **F**, Doxycycline (dox)-induced (1 μg/mL, 72 hours) TNIK shRNA reduces cell migration and invasion (Transwell assays) in SW900. Migrating and invading cells were stained with crystal violet and counted with ImageJ. Data represent mean ± SD from *n* = 3 independent experiments; one-way ANOVA, Tukey’s multiple comparisons post-test. **G**, Cell migration and invasion of SW900 after treatment with TNIK inhibitors (100 nM NCB-0846 or 1 μM INS18_055), or treatment with inactive control compound (100nM NCB-0970) for 72 hours. Migrating and invading cells were stained with crystal violet and counted with ImageJ. Data represent mean ± SD from *n* = 3 independent experiments; one-way ANOVA, Tukey’s multiple comparisons post-test.

To determine whether TNIK regulated pEMT in LUSC cells, we assessed whether depleting TNIK affected the expression of epithelial and mesenchymal markers in two LUSC models, SW900 and NCI-H520 cells, that exist in a hybrid E/M state. In these cells, TNIK depletion decreased expression of mRNAs encoding mesenchymal genes, including *VIM* (vimentin), *SNAI2* (SLUG), *TWIST2*, *TGFBI* (TGF-β induced), *TGFB1*, and *TCF4*, among others, and upregulated epithelial genes such as *CDH1* (E-cadherin), consistent with disruption of pEMT transcriptional programs (**Fig. 1B**). To determine whether these transcriptional changes translated into cell state reprogramming, TNIK was depleted for 72 and 144 hours, and the expression of E- and M-state markers was examined (**Fig. 1C**). TNIK loss consistently reduced expression of the M-state markers vimentin and SLUG in both SW900 and NCI-H520 cells at both time points, while E-cadherin protein levels increased (**Fig. 1C; Supplementary Fig. S1A, S1B**). These findings were further validated at single-cell resolution by fluorescence-activated cell sorting (FACS). FACS analysis showed that a large proportion of SW900 and NCI-H520 cells co-expressed both mesenchymal (vimentin or CD44) and epithelial (E-cadherin or EpCAM [Epithelial Cell Adhesion Molecule]), confirming that these LUSC cells exhibit a hybrid E/M state (**Fig 1D**; **Supplementary Fig. S1C**). Following TNIK knockdown, we observed a significant decrease in the expression (assessed as mean fluorescence intensity [MFI]) of the M-state-associated markers vimentin and CD44, in parallel with an increased MFI of E-state markers E-cadherin and EpCAM (**Fig. 1E; Supplementary Fig. S1D, S1E**). This increase in the expression of E-markers translated into a reduction of the fraction of hybrid E/M cells following TNIK knockdown (**Fig. 1D; Supplementary Fig. S1C**). Taken together, these results establish TNIK as a new regulator of pEMT in LUSC, and that its depletion reprograms hybrid E/M LUSC cells towards an epithelial state.

Given that TNIK depletion suppressed pEMT and promoted epithelial reprogramming, we next investigated whether TNIK-associated changes translated into altered pEMT-associated phenotypes. To this end, we determined the effects of TNIK depletion or pharmacologic inhibition on the migratory and invasive behavior of LUSC cells using Transwell migration and invasion assays. TNIK knockdown using doxycycline-inducible shRNAs significantly reduced the number of migrating and invading cells relative to control conditions, indicating impaired cell motility upon TNIK depletion in both SW900 and NCI-H520 cells (**Fig. 1F; Supplementary Fig. S1F**). Consistent with these findings, treatment of SW900 and NCI-H520 cells with TNIK inhibitors (the tool compound NCB-0846 [10] and the clinical-grade inhibitor INS18_055, also known as rentosertib [25, 26]) significantly reduced migratory and invasive capacities in both cell lines (**Fig. 1G; Supplementary Fig. S1G**). Importantly, treatment with NCB-0970, an inactive analog of NCB-0846 that does not inhibit TNIK [10], had no effects on cell migration or invasion (**Fig. 1G; Supplementary Fig. S1G**). Together, these results demonstrate that TNIK is a critical regulator of migratory and invasive behavior in LUSC cells and suggest that TNIK does so in a catalytic-dependent manner.

### TNIK maintains pEMT in LUSC by regulating MYC expression

The above findings indicated that TNIK is required to maintain a hybrid E/M state in LUSC cells and that its inhibition shifted M-state-associated traits toward the E state. We next investigated the mechanisms underlying epithelial reprogramming following TNIK targeting. Gene-set enrichment analysis (GSEA) of TNIK-depleted LUSC cells compared to controls identified candidate upstream regulators of pEMT affected by TNIK depletion, including Hippo, focal adhesion, cell cycle, pluripotency, TGF-β, and WNT/β-catenin signaling pathways, consistent with TNIK acting upstream of several plasticity-associated programs (**Fig. 2A**).

**Figure 2.**
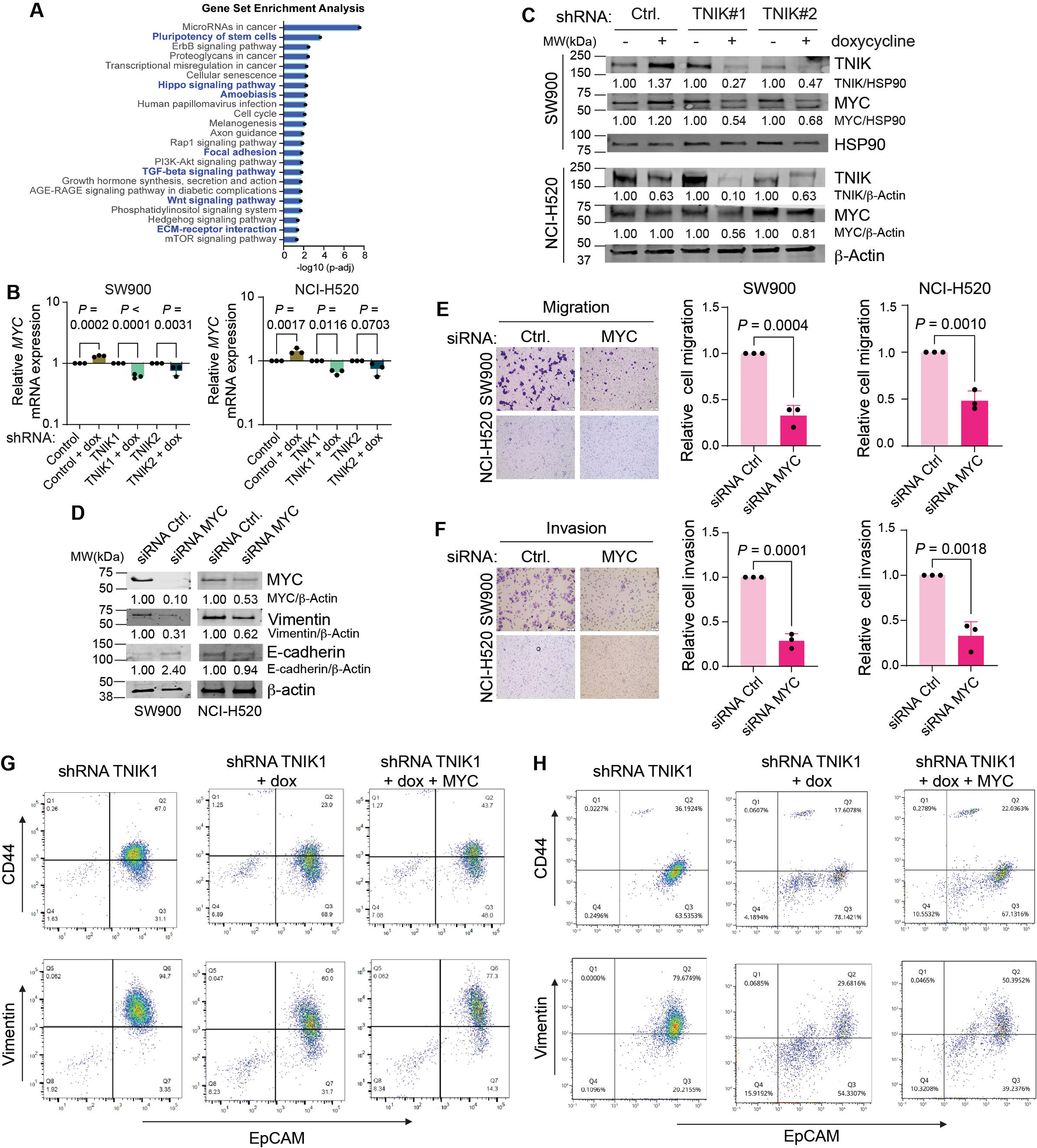
TNIK maintains hybrid E/M states in LUSC cells by regulating MYC expression. **A**, Identification of transcriptional programs regulated downstream of TNIK in LUSC cells. Gene Set Enrichment Analysis (GSEA) was performed on RNA-seq data from TNIK-depleted LUSC cells (SW900, NCI-H520, LK-2, and LC1/SQ). Top significantly downregulated pathways (*p-*adj shRNA *TNIK* vs shRNA control) are shown. Programs associate with pEMT are indicated in blue. **B**, RT-qPCR was performed to assess the expression of *MYC* and *RPL19* (internal control) in TNIK-depleted SW900 (left) and NCI-H520 (right) cells (shRNA induced with 1 μg/mL doxycycline (dox), 72 hours). RT-qPCR data were analyzed using the ΔΔCt method. Untreated (minus dox) samples were set as controls (relative target gene expression = 1.00). Data represent mean ± SD from *n* = 3 independent experiments in triplicate; one-way ANOVA, Tukey’s multiple comparisons post-test. **C**, Representative western blot analysis of TNIK and MYC levels in SW900 (top) and NCI-H520 (bottom) cells after 144 hours of TNIK knockdown (shRNA; 1 μg/mL dox replaced every 72 hours). HSP90 or β-actin were used as loading controls. Protein level (band intensity) relative to the respective loading control was normalized to untreated (minus dox) samples. The blots are representative of *n* = 3 independent experiments in each cell line. **D**, Western blot analysis of MYC, E-cadherin, and vimentin protein levels following MYC knockdown in SW900 (left) and NCI-H520 cells (right) with siRNA (50 nM, 72 hours). β-actin was used as a loading control. Protein level (band intensity) relative to β-actin was normalized to siRNA control samples. The blots are representative of *n* = 3 independent experiments in each cell line. **E, F**, Control or *MYC*-targeting siRNAs (50 nM, 72 hours) were transfected into SW900 and NCI-H520 cells. Transwell assays show reduced migration (E) and invasion (F) after MYC knockdown in SW900 and NCI-H520 cells. Migrating and invading cells were stained with crystal violet and counted with ImageJ. Data represent mean ± SD from *n* = 3 independent experiments; two-tailed *t* test. **G, H**, Representative histograms of EpCAM, vimentin, and CD44 expression by flow cytometry in TNIK-depleted SW900 (G) and NCI-H520 (H) after dox-induced shRNA (1 μg/mL, 144 hours total treatment, replaced every 72 hours) followed by expression of MYC- or -empty vector (72 hours). The histogram is representative of *n* = 3 independent experiments.

TNIK has previously been implicated in the WNT/β-catenin signaling pathway, where it promotes transcription of downstream targets. Additional studies have shown that TNIK regulates the Hippo effectors YAP/TAZ [8, 10, 13–15]. Given that MYC is a canonical target of both the WNT/β-catenin and YAP/TAZ pathways, and a central regulator of cell-cycle progression, pluripotency, and pEMT [10, 27–31], we examined MYC as a candidate effector linking TNIK to pEMT.

TNIK depletion resulted in loss of MYC expression at both the mRNA and protein levels (**Fig. 2B** and **C**). This effect was recapitulated pharmacologically, whereby treatment of SW900 or NCI-H520 cells with the TNIK inhibitor NCB-0846 reduced total and nuclear MYC protein levels, whereas the inactive control compound, NCB-0970, did not affect MYC expression (**Supplementary Fig. S2A and S2B**). TNIK inhibition also reduced the pEMT-associated transcription factor SOX9, a WNT/β-catenin target [32–34] (**Supplementary Fig. S2C**), further supporting suppression of WNT-associated transcription downstream of TNIK loss. Together, these data identify MYC as a key transcriptional target through which TNIK sustains the hybrid E/M state in LUSC.

To test whether TNIK regulated pEMT through MYC, first, we silenced MYC using siRNA in SW900 and NCI-H520 cells (**Fig. 2D**). MYC depletion decreased vimentin expression in both cell lines and increased E-cadherin expression in SW900 cells (**Fig. 2D**). Functionally, MYC silencing significantly reduced migration and invasion through Matrigel-derived extracellular matrix, as assessed in Transwell assays (**Fig. 2E** and **F**). Therefore, depleting MYC functionally reprogrammed LUSC cells towards an epithelial state and phenocopied the effects of TNIK loss.

Next, we evaluated whether re-expression of MYC would restore mesenchymal traits in TNIK-depleted cells (**Supplementary Fig. S2D**). FACS analysis for epithelial and mesenchymal markers showed that re-expressing MYC reversed the effects of TNIK depletion; MYC expression in TNIK-depleted cells led to increased levels of mesenchymal markers (vimentin and CD44), decreased expression of the epithelial marker EpCAM, and restoration of the hybrid E/M state (**Fig. 2G and H**; **Supplementary Fig. S2E, S2F and S2G**). Collectively, these results identify MYC as a central downstream effector through which TNIK maintains a hybrid E/M state in LUSC cells.

### TNIK-loss-mediated epithelial reprogramming is associated with induction of a MYC-dependent senescent-like state

The results above indicate that TNIK loss reprogrammed LUSC cells towards an epithelial state by downmodulating MYC. To further explore the relevance of TNIK and MYC interactions in LUSC, we next investigated whether TNIK-loss-mediated epithelial reprogramming affected cell proliferation, as mesenchymal-to-epithelial reprogramming has been associated with increased proliferation [35–38]. We assessed cell-cycle phase distribution in control and TNIK-depleted SW900 and NCI-H520 cells using 5-ethynyl-2’-deoxyuridine (EdU) incorporation and DNA staining. TNIK depletion significantly reduced the proportion of EdU-positive cells, indicative of reduced DNA synthesis (**Fig. 3A and B; Supplementary Fig. S3A and S3B**). Cell-cycle phase analysis revealed a significant decrease in the fraction of cells in S phase, consistent with the reduction of EdU-positive cells, as well as a significant increase in the fraction of cells in sub-G0/G1 (dead) (**Fig. 3A and B; Supplementary Fig. S3A and S3B**). These results are compatible with reduced proliferative capacity of LUSC cells following TNIK depletion. Next, we evaluated the effects of pharmacologic inhibition of TNIK on long-term cell proliferation. Colony formation assays in SW900 and NCI-H520 cells treated with increasing concentrations of TNIK inhibitors NCB-0846 or INS18_055 showed a significant reduction in colony-forming capacity at doses known to selectively target TNIK activity [10, 25], with NCI-H520 cells being more sensitive to TNIK inhibition (**Fig. 3C and D**). Importantly, the inactive compound NCB-0970 had minor effects on SW900 or NCI-H520 cell proliferation (**Fig. 3C and D**).

**Figure 3.**
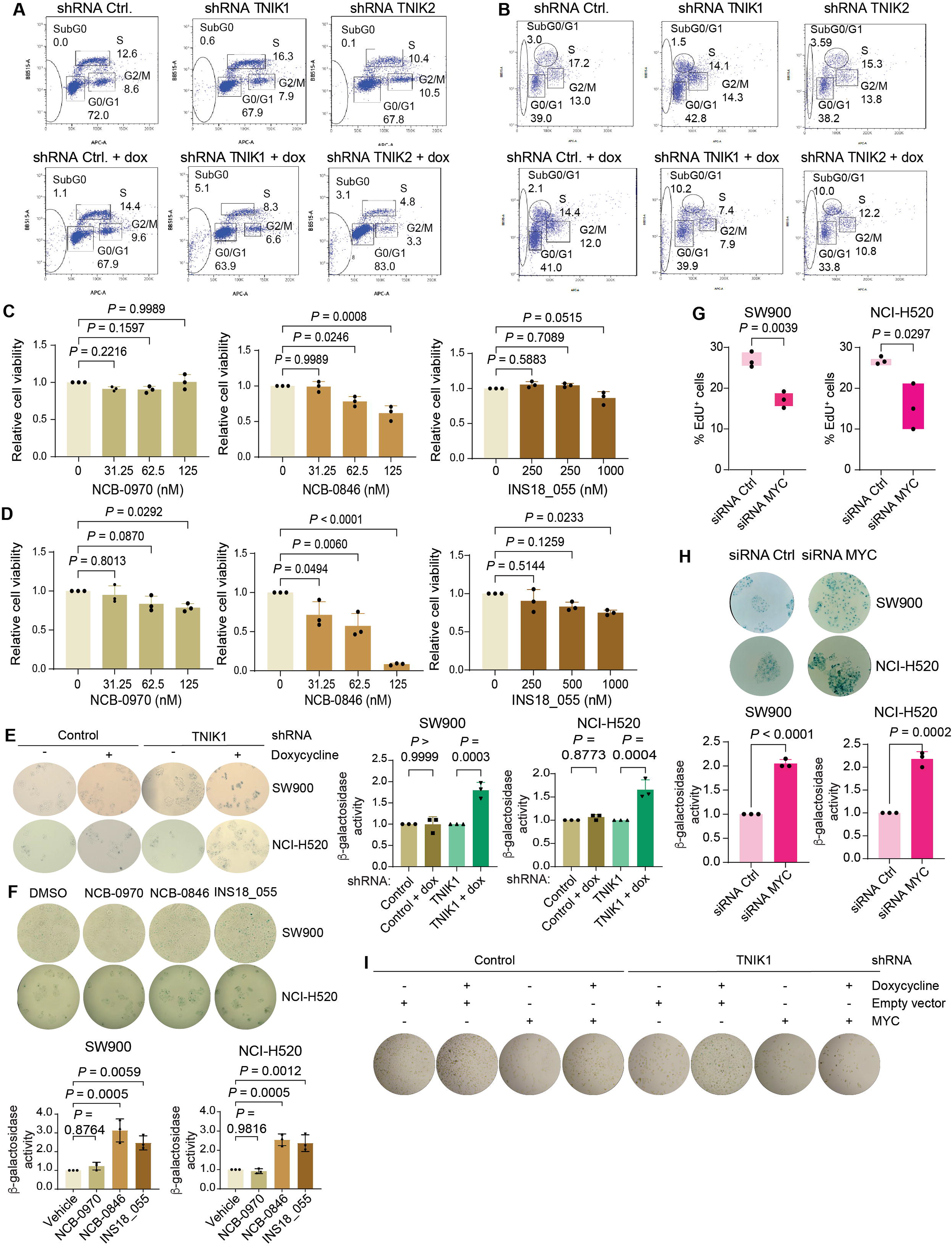
Targeting TNIK suppressed LUSC cell proliferation and induced a MYC-dependent senescence-like phenotype. **A, B**, Representative cell cycle distribution analysis of SW900 (A) and NCI-H520 (B) cells after doxycycline-inducible shRNA *TNIK* knockdown (1 μg/mL, 96 hours). Cells were labeled with EdU (10 μM; 1 h), harvested, and fixed. DNA content and EdU incorporation were analyzed by flow cytometry. Histograms are representative of *n* = 3 independent experiments. **C, D**, Colony-forming capacity (12 days) of SW900 (C) and NCI-H520 (D) cells after treatment with NCB-0970, NCB-0846, or INS18_055 at the concentrations indicated, with inhibitors replaced every 72-h. Data represent mean ± SD from *n* = 3 independent experiments in triplicate; one-way ANOVA, Tukey multiple comparisons post-test. **E**, *Top*, senescence-associated β-galactosidase (SA-β-gal) activity was examined in control or TNIK-depleted (1 μg/mL dox-induced shRNA, 144 hours) SW900 and NCI-H520 cells. Cells were imaged after incubation with X-gal solution, and senescent cells were identified by blue cytoplasmic staining. *Bottom,* quantification of SA-β-gal-positive cells was performed using ImageJ and normalized to untreated (minus dox) controls. Data represent mean ± SD from *n* = 3 independent experiments with triplicates; one-way ANOVA, Tukey multiple comparisons post-test. **F**, *Top*, SA-β-gal activity was examined in SW900 and NCI-H520 cells after treatment with NCB-0970 (100 nM, inactive control compound for NCB-0846), NCB-0846 (100 nM), or INS18_055 (1000 nM) for 72 hours. *Bottom,* quantification of SA-β-gal-positive cells was performed using ImageJ and normalized to the DMSO-treated control. Data represent mean ± SD from *n* = 3 independent experiments with triplicates. One-way ANOVA, Tukey multiple comparisons post-test. **G**, Control or *MYC*-targeting siRNAs (50 nM, 72 hours) were transfected into SW900 and NCI-H520 cells. EdU-positive cells were quantified by FACS analysis. Data represent mean ± SD from *n* = 3 independent experiments; two-tailed *t-*test. **H**, *Top*, SA-β-gal activity was examined in control or MYC-depleted SW900 and NCI-H520 cells (siRNA 50 nM; 72 hours). Cells were imaged after incubation with X-gal solution. *Bottom,* senescent cells were identified by blue cytoplasmic staining using ImageJ. Data represent mean ± SD from *n* = 3 independent experiments with triplicates; two-tailed *t*-test. **I**, SA-β-gal activity was examined in control or TNIK-depleted SW900 cells transfected with a MYC-expressing or empty vector. Senescent cells were identified by blue cytoplasmic staining following incubation with X-gal solution.

To corroborate whether the observed reduction in EdU incorporation was associated with cellular senescence, as suggested by our RNA-seq analyses, we assessed senescence-associated β-galactosidase (SA-β-gal) activity. TNIK depletion significantly increased SA-β-gal activity in both SW900 and NCI-H520 cells (**Fig. 3E; Supplementary Fig. S3C**). This effect was accompanied by an approximate 2-fold increase in the expression of senescence-associated secretory factors *IL6* and *IL8* (**Supplementary Fig. S3D**). Similarly, treatment with TNIK inhibitors NCB-0846 and INS18_055 significantly increased the fraction of β-gal-positive cells in both cell lines, whereas NCB-0970 showed no effect on β-gal activity (**Fig. 3F**), suggesting that TNIK activity is required to prevent senescence in LUSC cancer cells.

Lastly, we investigated whether MYC contributed to TNIK-dependent regulation of cell proliferation and senescence. Silencing of MYC (siMYC) in SW900 and NCI-H520 cells resulted in a significant reduction of EdU-positive cells and a significant increase in the proportion of β-gal-positive cells (**Fig. 3G and H**), indicating that MYC loss phenocopied the effects of TNIK depletion on LUSC cell proliferation and senescence. To corroborate whether reduced MYC levels were responsible for TNIK inhibition-induced senescence regulation, we re-expressed MYC in TNIK-depleted cells. Whereas TNIK knockdown led to increased SA-β-gal activity, restoring MYC reversed this effect (**Fig. 3I**; **Supplementary Fig. S3E)**. Together, our data are consistent with TNIK depletion inducing an epithelial, low-proliferative state in LUSC in a MYC-dependent manner.

### TNIK and EZH2 co-inhibition suppressed pEMT and activated stress signaling in LUSC cells

Our data indicate that TNIK couples epithelial-mesenchymal plasticity and cell proliferation in LUSC cells. To explore candidate therapeutic strategies involving TNIK inhibitors, we performed a small-molecule screen of 75 clinically relevant cancer compounds (**Supplementary Table 2**). NCI-H520 cells were treated with each compound in the presence or absence of the TNIK inhibitor NCB-0846 (EC10). The screen identified two chemotherapeutic agents (5-fluorouracil and oxaliplatin) that cooperated with TNIK inhibition, suggesting that TNIK inhibition may enhance responses to platinum-based chemotherapy, a first-line LUSC treatment [2] (**Fig. 4A**). This screen also identified two epigenetic inhibitors, (+)-JQ1 and tazemetostat, that were of further interest, as both BET proteins and EZH2 have been reported to cooperate with MYC in the transcriptional regulation of cell proliferation and survival [29, 39] (**Fig. 4A**).

**Figure 4.**
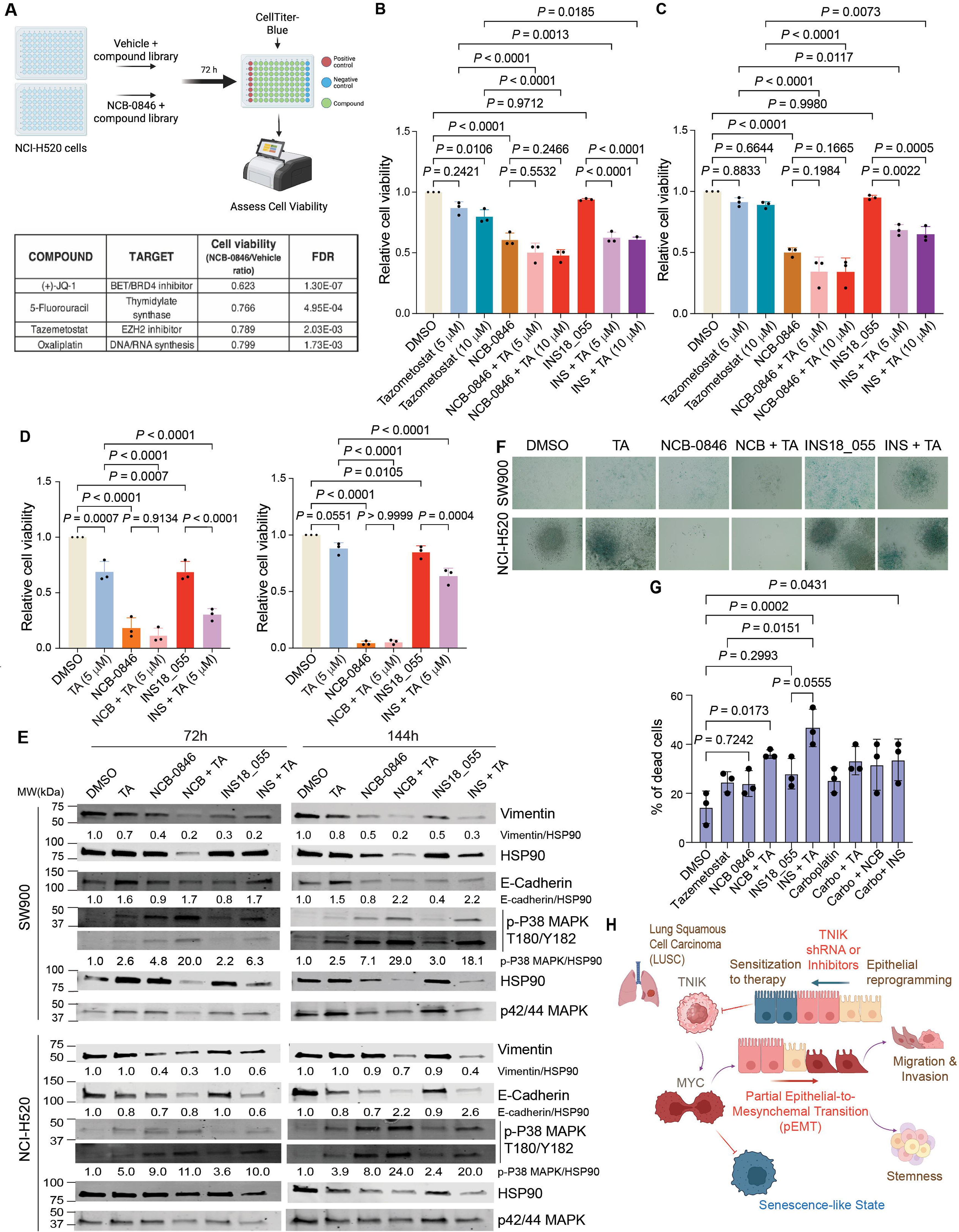
TNIK and EZH2 co-inhibition suppressed pEMT and activated stress signaling in LUSC cells. **A**, To identify therapeutic combinations that enhance TNIK inhibition, a library of clinically relevant small-molecule inhibitors was screened (at 100 nM and 1000 nM) in NCI-H520 treated with DMSO or NCB-0846 (at EC10). Hits (FDR < 5%) were prioritized based on biological activity (> 15% increase in sensitization to NCB-0846). The schematic was generated with BioRender. **B, C**, Cell viability (crystal violet assay) of SW900 (B) and NCI-H520 cells (C) treated with TNIK inhibitors (100 nM NCB-0846 or 1 μM INS18_055), or EZH2 inhibitor tazemetostat (TA; 5 or 10 µM), alone or in combination for 72 hours. Data represent mean ± SD from *n* = 3 independent experiments in triplicate; one-way ANOVA, Tukey’s multiple comparisons post-test. **D**, Cell viability (crystal violet assay) of SW900 (left) or NCI-H520 cells (right) treated with TNIK inhibitors (100 nM NCB-0846 or 1 μM INS18_055), or EZH2 inhibitor tazemetostat (TA; 5 µM) for 144 hours (inhibitors replaced every 72 hours). Data represent mean ± SD from *n* = 3 independent experiments in triplicate; one-way ANOVA, Tukey’s multiple comparisons post-test. **E**, pEMT marker expression (*n* = 3 independent experiments) was analyzed by immunoblot in SW900 (top) and NCI-H520 cells (bottom) treated with the indicated inhibitors (100 nM NCB-0846 [NCB], 1 μM INS18_055 [INS], or 5 µM tazemetostat [TA]) alone or in combination (72 and 144 hours). HSP90 and p42/p44 MAPK were used as loading controls. Protein level (band intensity) relative to HSP90 was normalized to DMSO-treated (control) samples. **F**, Senescence-associated (SA)-β-galactosidase activity was examined in SW900 (top) and NCI-H520 cells (bottom) treated as in D for 14 days (inhibitors were replaced every 72 hours). Images are representative of *n* = 3 independent experiments. **G**, LUSC organoids (XDO-4242) were treated (48 h) with single-agent TNIK inhibitors (250 nM NCB-0846 or 1 µM INS18_055), Tazemetostat (TA, 5 µM), carboplatin (6 µM), or the indicated combinations. Organoids were then collected, dissociated, and dead cells were quantified using a trypan blue exclusion assay. Data represent mean ± SD from *n* = 3 independent experiments; one-way ANOVA, Tukey’s multiple comparisons post-test. **H**, Proposed model of how TNIK contributes to pEMT and evasion of senescence by regulating MYC expression in LUSC cells. Conversely, inhibiting TNIK reprograms LUSC cells towards an epithelial, senescent-like state. The schematic was generated with BioRender.

To verify the screening results, we first investigated whether inhibiting TNIK would sensitize LUSC cells to standard of care platinum-based chemotherapy. We investigated two strategies: combination treatment, in which cells were simultaneously treated with TNIK inhibitors and carboplatin, or sequential treatment, in which cells were first treated with carboplatin, followed by TNIK inhibitors (NCB-0846 or INS18_055). Simultaneous treatment with carboplatin and TNIK inhibitors was not significantly stronger than single treatment with TNIK inhibitors (**Supplementary Fig. S4A and S4B**), whereas combined treatment with cisplatin and TNIK inhibitor was significantly stronger than either single treatment in NCI-H520 cells (*P* = 0.0267 for NCB-0846 treatment versus NCB-0846 + cisplatin combination, *P* = 0.0250 for cisplatin treatment versus NCB-0846 + cisplatin combination; **Supplementary Fig. S4B**). However, sequential treatment of SW900 cells with carboplatin, followed by treatment with NCB-0846, was significantly more effective than NCB-0846 alone (*P =* 0.035 for NCB-0846 treatment versus sequential carboplatin + NCB-0846 combination), and showed a trend towards improvement over carboplatin alone (*P =* 0.0585 for carboplatin treatment versus sequential carboplatin + NCB-0846 combination; **Supplementary Fig. S4A**).

Next, we investigated combinations of TNIK inhibitors (NCB-0846 and INS18_055) with tazemetostat, given the role of EZH2 in cell plasticity and its known functional interactions with MYC [39–41]. Further, inhibition of EZH2 has been proposed as a strategy to treat SOX2-associated cancers; SOX2 is a stemness and cell-plasticity transcription factor that is generally co-amplified with *TNIK* in LUSC as part of the common 3q amplicon [42–44].

Short-term treatment (72 hours) of SW900 and NCI-H520 cells with the combination of tazemetostat and the clinical-grade TNIK inhibitor INS18_055 (each at EC10) significantly decreased cell viability in both cell lines compared with single treatment (SW900 cells: *P =* 0.0013 tazemetostat versus combination and *P <* 0.0001 INS18_055 versus combination; NCI-H520 cells: *P =* 0.0117 tazemetostat versus combination and *P =* 0.0022 INS18_055 versus combination; **Fig. 4B and C**). The effects of tazemetostat and INS18_055 combination were further amplified after treatment for 144 hours, with inhibitors replaced every 72 hours (SW900 cells: ∼40% reduction of cell viability with single treatment tazemetostat or INS18_055, *P =* 0.0007 tazemetostat or INS18_055 versus vehicle; ∼70% reduction in cell viability with combined tazemetostat and INS18_055, *P* < 0.0001 tazemetostat or INS18_055 alone versus combination; NCI-H520 cells: ∼10% reduction of cell viability with single treatment tazemetostat, *P =* 0.0551 tazemetostat versus vehicle; ∼15% reduction of cell viability with single treatment INS18_055, *P =* 0.0105; ∼ 40% reduction of cell viability with combined tazemetostat and INS18_055, *P* < 0.0001 tazemetostat alone versus combination, *P =* 0.0004 INS18_055 alone versus combination; **Fig. 4D**). Together these data support that EZH2 inhibition cooperates with TNIK catalytic inhibitors to reduce LUSC cell viability.

We next tested whether the cooperation between TNIK and EZH2 inhibition involved convergent suppression of pEMT. Immunoblot analysis revealed that single-treatment with TNIK inhibitors, NCB-0846 or INS18_055 (72 and 144 hours), reduced the expression of the mesenchymal marker vimentin by ∼50%, consistent with loss of mesenchymal traits after targeting TNIK (**Fig. 4E**). Tazemetostat had modest effects on the expression of vimentin but increased the expression of E-cadherin by ∼1.5-fold only in SW900 cells (**Fig. 4E**). The combination of TNIK inhibitors with tazemetostat further reduced the expression of vimentin by ∼70% and caused a 2-fold-increase in the expression of E-cadherin in both SW900 and NCI-H520 cells (**Fig. 4E**).

We also investigated whether EZH2 inhibition with tazemetostat increased the senescent-like phenotype triggered by TNIK inhibitors. Inhibition of TNIK alone increased the fraction of β-gal-positive cells (**Fig. 4F**, consistent with Fig. 2). However, treatment with tazemetostat did not increase β-gal activity in TNIK-inhibited cells (**Fig. 4F**), indicating that the reduced cell viability of the combination is not explained by augmented senescence.

Because the hybrid E/M state is thought to confer adaptability and stress tolerance [20], we reasoned that the more pronounced suppression of pEMT achieved by combined TNIK and EZH2 inhibition might instead render cells more susceptible to therapeutic stress. Consistent with this, immunoblot analysis of p38 MAPK, a stress-activated kinase that couples cellular stress to growth-suppressive programs and can be regulated by TNIK-related kinases [45, 46], showed a 2- to 5-fold increase in the activation of p38 MAPK (assessed by phosphorylation at T180/Y182) after treatment with TNIK inhibitors alone in both SW900 and NCI-H520 cells (**Fig. 4E**). Activation of p38 MAPK was further amplified (by ∼5-fold over TNIK inhibitors alone) when cells were co-treated with TNIK inhibitors and tazemetostat (**Fig. 4E**), indicating that beyond its effects on suppressing pEMT, the combination of TNIK inhibitors with tazemetostat engaged stress-response signaling.

Last, we evaluated treatment responses in a LUSC patient-derived organoid model (XDO-4242 [47]). Organoids were treated for 48 h with TNIK inhibitors, tazemetostat, carboplatin, or combinations thereof. Consistent with observations in LUSC cell-line models, the combination of TNIK inhibitors and tazemetostat resulted in a significant increase in the fraction of dead cells (36% dead cells; *P =* 0.0173 for the combination of NCB-0846 + tazemetostat versus DMSO vehicle control; 46% dead cells; *P =* 0.0002 for the combination of INS18_055 + tazemetostat versus control), demonstrating enhanced cytotoxic effects of the combination (**Fig. 4G**). The combination of INS18_055 with carboplatin similarly increased the proportion of dead cells in the XDO-4242 organoid model (34% dead cells; *P =* 0.0431 versus control).

In closing, our results suggest that TNIK sustains a MYC-driven pEMT state that coupled cellular plasticity with proliferation and evasion of senescence, and reveal that its inhibition, alone or in combination with EZH2 inhibitors, represents a potential strategy in this subset of LUSC tumors.

## DISCUSSION

Partial EMT has been gathering attention as a major contributor to tumor progression and therapy resistance in squamous cell carcinomas, including LUSC [21, 48–51]. Here, we show that the serine/threonine protein kinase TNIK, which is recurrently amplified in LUSC, acts as a coordinator of pEMT and cell proliferation through the regulation of the MYC oncogene.

Our study revealed that depleting or inhibiting TNIK reprogrammed LUSC cells towards an epithelial, non-proliferative state by reducing the expression of the oncogenic transcription factor MYC (**Fig. 4H**). Transcriptomic analyses of TNIK-depleted LUSC cells position TNIK upstream of multiple transcriptional programs that promote pEMT, including the TGF-β/SMAD, WNT/ β-catenin and YAP/TAZ programs [23, 49–52]. Consistent with these transcriptional changes, TNIK depletion or inhibition reduced the expression of mesenchymal markers and resulted in impaired cell migration and invasion. Mechanistically, we identified MYC as a key downstream effector of TNIK in sustaining the hybrid E/M state in LUSC. We found that TNIK depletion reduced *MYC* mRNA expression, MYC knockdown recapitulated key aspects of TNIK loss, and restoring MYC expression reversed the epithelial reprogramming induced by TNIK depletion. Consistent with a role for TNIK in maintaining mesenchymal and plasticity-associated programs, TNIK inhibition with NCB-0846, but not its inactive analog NCB-0970, has been shown to suppress mesenchymal regulators such as SLUG and SNAIL in colorectal cancer models [10], supporting a broad role for TNIK in sustaining epithelial–mesenchymal plasticity across cancer types.

While mesenchymal-to-epithelial reprogramming can lead to increased cancer cell proliferation [35–37], we found that TNIK depletion or inhibition decreased cell proliferation, as assessed by reduced EdU incorporation, a surrogate measure of DNA synthesis, and triggered a senescent-like state characterized by enhanced SA-β-gal activity and expression of senescence-associated secretory factors IL6 and IL8. Notably, up to 80% of LUSC tumors are *TP53*-mutant (upstream of p21) and 30% are *CDKN2A*-deleted (encoding p16) [53], suggesting that these tumors may trigger a non-canonical senescence program. Our data indicate that the effects of TNIK on senescent traits were similarly MYC-dependent, as restoring MYC levels prevented the SA-β-gal activity in TNIK-depleted SW900 cells. These findings are consistent with reports that MYC loss promotes senescence whereas sustained MYC expression supports proliferative capacity [54, 55], providing new insights into how TNIK contributes to tumorigenesis by coordinating pEMT and cell proliferation.

The fact that TNIK inhibition suppressed pEMT and induced a senescence-like state in LUSC cells may also have therapeutic implications for the use of TNIK inhibitors in LUSC, which are currently entering Phase III clinical trials for idiopathic pulmonary fibrosis [25, 26]. Our small-molecule screening to find cooperative partners of TNIK inhibitors identified platinum-based chemotherapeutics, which are standard of care in LUSC [2]. However, we did not observe significant benefit of co-treatment with NCB-0846 or INS18_055 with carboplatin or cisplatin in LUSC cells, consistent with TNIK inhibition reducing DNA synthesis (as indicated by EdU incorporation assays). Instead, we found positive activity when TNIK inhibitors were provided after carboplatin, consistent with prior reports whereby inhibiting TNIK reduced cell viability in platinum-resistant ovarian cancer [13]. These observations indicate that the interaction between TNIK inhibition and platinum-based chemotherapeutics is agent- and schedule-dependent and suggest that inhibiting TNIK may be a strategy in specific contexts to target potential persister and platinum-resistant cells. Indeed, drug-tolerant persister cells share transcriptional and functional similarities with pEMT states, including stress tolerance and tumor-initiating potential, which contribute to treatment resistance and tumor relapse [20, 22, 56–58]. In this regard, we found that when TNIK inhibitors were combined with the methyltransferase EZH2 inhibitor tazemetostat, identified in our small-molecule screening, cell viability was significantly reduced without increasing SA-β-gal activity, although the combination further suppressed pEMT. Instead, co-treatment with TNIK and EZH2 inhibitors triggered a therapy-induced stress response characterized by activation of p38 MAPK, consistent with the loss of an adaptive hybrid E/M state.

Lastly, our results provide a rationale for whether inhibiting TNIK could affect LUSC responses to immune checkpoint blockade (ICB) by reversing pEMT and/or promoting acute senescence. Whereas pEMT is known to promote resistance to ICB [59, 60], the connection between senescence and ICB is more complex, with reports of senescence promoting or preventing responses to ICB. While senescent tumor cells are initially immunogenic and can be eliminated by NK and CD8+ T cells [61, 62], the accumulation of senescent cells may promote an immunosuppressive tumor microenvironment [63–65]. In this regard, a recent study in small cell lung cancer (SCLC) by Tanimoto and colleagues [34] identified TNIK as a dependency in the SCLC-P subtype, which is characterized by high MYC expression. Mechanistically, TNIK inhibition with NCB-0846 reduced MYC and SOX9 levels in SCLC cells and sensitized these cells to anti-PD-L1 blockade *in vivo*. Importantly, this effect was mediated by changes in the cancer cell secretome, primarily CCL2. However, whether inhibiting TNIK in MYC-high SCLC cells triggered senescence or affected cell plasticity was not determined. Nonetheless, together with our findings, these data support a model in which MYC represents a conserved downstream dependency of TNIK signaling across lung cancer subtypes, and likely across multiple solid malignancies [10, 13, 34].

Collectively, these results implicate TNIK in the mechanisms linking epithelial-mesenchymal plasticity with proliferation and evasion of senescence via regulation of the MYC oncogene, and provide additional mechanistic insights into therapeutic strategies for the clinical deployment of TNIK inhibitors in LUSC and other TNIK-dependent malignancies.

## AUTHOR CONTRIBUTIONS

MH was responsible for most of the experimental work and helped conceptualize the study. MH, KO, KWH, and SS conducted experiments and analyzed the data. YZ conducted bioinformatic analyses. PT-A developed the concept, acquired funding, and supervised the overall course of the experiments. MH and PT-A wrote the manuscript. All authors reviewed and edited the manuscript.

## ACKNOWLEDGEMENTS

We thank members of the Torres-Ayuso lab, and Dr. Erica Golemis for helpful discussions. We acknowledge Drs. Mary Barbe (Lewis Katz School of Medicine), Margret Einarson (Fox Chase Cancer Center), and Amir Yahmarmoodi (Lewis Katz School of Medicine) for technical assistance with microscopy, small-molecule screening, and flow cytometry, respectively. We thank Drs. Ming Tsao, Nikolina Radulovich, and Molly Udaskin (University Health Network, Toronto, ON, Canada) for providing technical advice for maintaining LUSC organoids.

## ETHICS DECLARATIONS

### Ethics

No studies involving human participants or patients were performed in this study.

## DATA AVAILABILITY STATEMENT

The data generated in this study are available within the article and its supplementary data files and can be made available upon request to Dr. Pedro Torres-Ayuso. The sequencing data generated in this study are publicly available in Gene Expression Omnibus (GEO) at GSE345484.

