## Supplementary Figures, Tables, and Methods for "TNIK maintains a MYC-driven partial EMT state that supports proliferation and evasion of senescence in lung squamous cell carcinoma"

**Supplementary Figures and Supplementary Figure Legends; Supplementary Tables; Supplementary Methods and Supplementary References.**

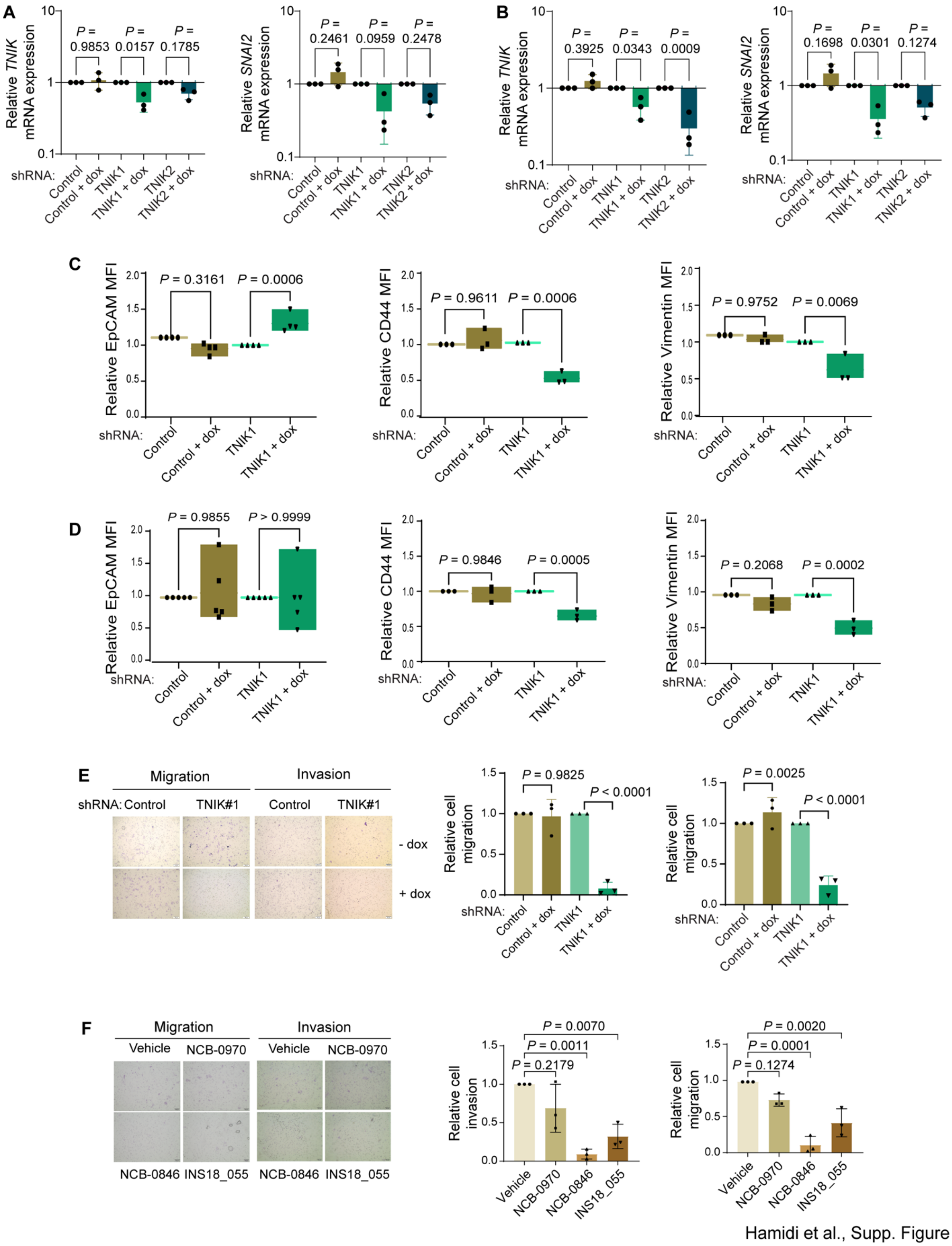


Hamidi et al., Supp. Figure 1

**Supplementary Figure S1. TNIK depletion reprograms LUSC cancer cells towards an epithelial cell state.**

**A, B,** RT-qPCR analysis of *TNIK*, *SNAI2,* and *RPL19* (internal control) expression after shRNA-mediated *TNIK* depletion (doxycycline (dox)-induced, 1 µg/mL, 72 h) in SW900 (A) and NCI-520 (B) cells. RT-qPCR data were analyzed using the ΔΔCt method; untreated (minus dox) samples were set as controls (relative target gene expression = 1.00). Data represent mean ± SD from *n* = 3 independent experiments in triplicate; one-way ANOVA, Tukey's multiple comparisons post-test.

**C, D,** Flow cytometry analysis of mean-fluorescence intensity (MFI) value of vimentin, CD44, EpCAM, in SW900 (C) and NCI-H520 (D) cells treated after dox-induced TNIK knockdown. Analysis was performed using FlowJo software. Data represent mean ± SD from *n* = 3 independent experiments for SW900 cells, and 5 independent experiments for NCI-H520 cells.

**E,** Dox-induced (1 µg/mL, 72 h) *TNIK* shRNA reduces cell migration and invasion (Transwell assays) in NCI-H520 cells. Migrating and invading cells were stained with crystal violet and counted with ImageJ. Data represent mean ± SD from *n* = 3 independent experiments; one-way ANOVA, Tukey's multiple comparisons post-test.

**F**, Cell migration and invasion of NCI-H520 cells after treatment with TNIK inhibitors (100 nM NCB-0846 or 1 µM INS18_055), or treatment with inactive control compound (100 nM NCB-0970) for 72 hours. Migrating and invading cells were stained with crystal violet and counted with ImageJ. Data represent mean ± SD from *n* = 3 independent experiments; one-way ANOVA, Tukey's multiple comparisons post-test.

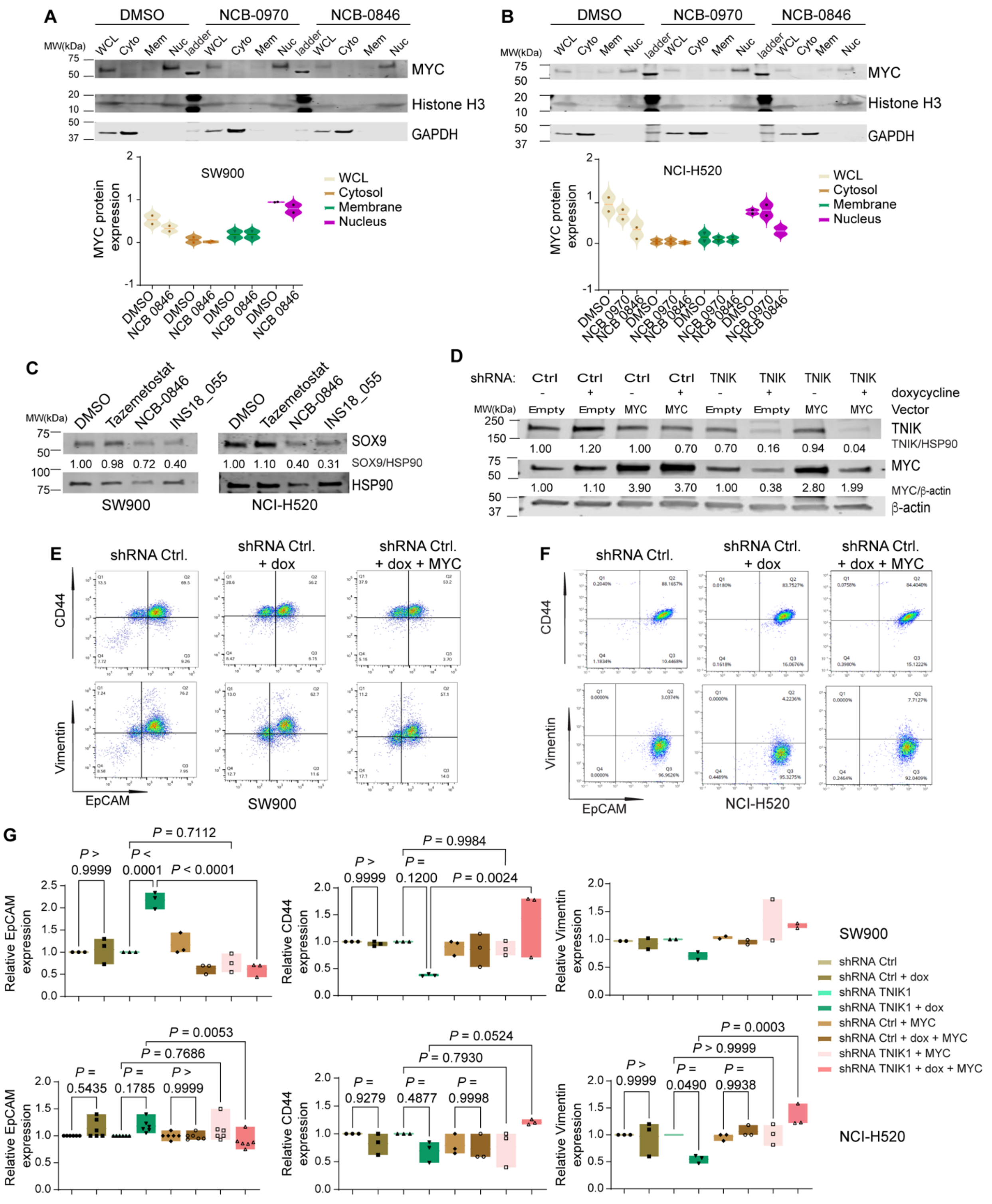


Hamidi et al., Supp. Figure 2

**Supplementary Figure S2. TNIK maintains pEMT in LUSC by regulating MYC expression.**

**A, B,** SW900 (A) and NCI-H520 (B) cells were treated for 72 hours with vehicle (DMSO) or TNIK inhibitor NCB-0846 (100 nM) or control inactive compound, NCB-0970 followed by cell fractionation to analyze MYC nuclear (Nuc), membrane (Mem), and cytosolic (Cyto) protein distribution. GAPDH and Histone H3 were used as cytosolic or nuclear fraction markers and loading controls, respectively. Protein level (band intensity) relative to fraction-specific control, or relative to GAPDH for whole-cell lysates (WCL) was normalized to vehicle-treated samples (control). The blot is representative of *n* = 2 independent experiments.

**C,** Western blot analysis of SOX9 in whole-cell extracts from SW900 (left) or NCI-H520 (right) cells treated with TNIK inhibitors (100 nM NCB-0846 or 1000 nM INS18_055) or inactive control compound NCB-0970 (100 nM) for 144 h. HSP90 was used as a loading control. Protein level (band intensity) relative to HSP90 was normalized to DMSO-treated samples. The blot is representative of *n* = 3 independent experiments.

**D,** Representative Western blot of TNIK and MYC expression in TNIK-depleted SW900 and NCI-H520 (1 µg/mL dox-induced shRNA, 144 h, replaced every 72 h) followed by expression of MYC- or -empty vector (72 h). Protein levels (band intensity) relative to β-actin were normalized to untreated samples. The blot is representative of *n* = 3 independent experiments.

**E, F,** Representative histograms of EpCAM, vimentin, and CD44 expression by flow cytometry in SW900 (E) and NCI-H520 (F) (1 µg/mL dox-induced shCtrl, 144 h, replaced every 72 h) followed by expression of MYC- or -empty vector (72 h). The histogram is representative of *n* = 3 independent experiments.

**G,** Representative quantification of CD44, EPCAM, and vimentin-positive cells of SW900 and NCI-H520 (bottom). Quantification was performed using FlowJo software, and data are expressed as a percentage of cells. Data represent mean ± SD from n = 6 independent experiments for EpCAM in the NCI-H520 cell line, and n = 3 independent experiments for the remaining experiments. Statistical significance was determined using one-way ANOVA, with Tukey's multiple comparisons post-test.

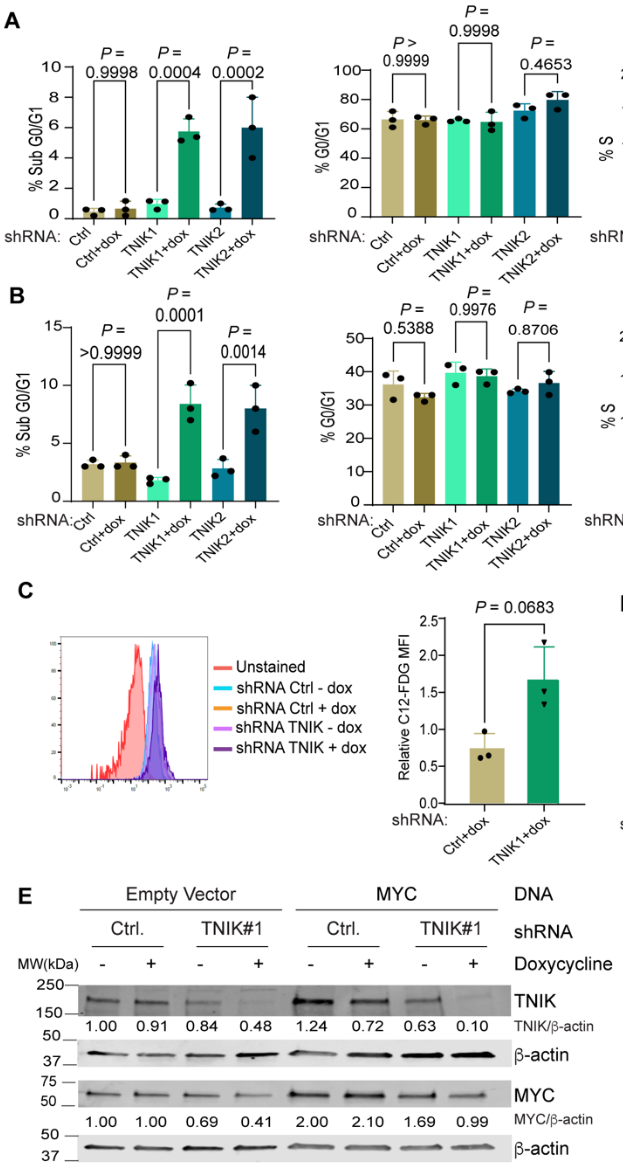


Hamidi et al., Supp. Figure 3

**Supplementary Figure S3. TNIK depletion or inhibition suppressed cell proliferation and triggered a senescence-like phenotype.**

**A, B,** Cell cycle phase distribution of SW900 (A) and NCI-H520 (B) cells after doxycycline-induced TNIK knockdown (1 μg/mL, 96 h) was performed using FlowJo software, and data represent mean ± SD of the percentage of cells on each phase (sub-G0/G1, G0/G1, S, and G2/M). *n* = 3 independent experiments; one-way ANOVA, Tukey's multiple comparisons post-test.

**C,** Senescence was assessed using C12-FDG staining (fluorescent β-galactosidase substrate) following depletion of TNIK (1 μg/mL dox-induced, 72 h) in SW900 cells. The histogram is representative of *n* = 3 independent experiments. Data represent mean ± SD from *n* = 3 independent experiments; two-tailed *t*-test.

**D,** RT-qPCR analysis of *IL-6, IL-8,* and *RPL19* (internal control) expression after shRNA-mediated *TNIK* depletion (1 μg/mL dox-induced, 72 h) in SW900 cells. RT-qPCR data were analyzed using the ΔΔCt method; untreated (minus dox) samples were set as controls (relative target gene expression = 1.00). Data represent mean ± SD from *n* = 3 independent experiments in triplicate; one-way ANOVA, Tukey's multiple comparisons post-test.

**E,** Representative Western blot of TNIK and MYC expression in TNIK-depleted SW900 and NCI-H520 (1 μg/mL dox-induced shRNA) concurrent with expression of MYC- or -empty vector (72 h). Protein levels (band intensity) relative to β-actin were normalized to untreated samples. The blot is representative of *n* = 3 independent experiments.

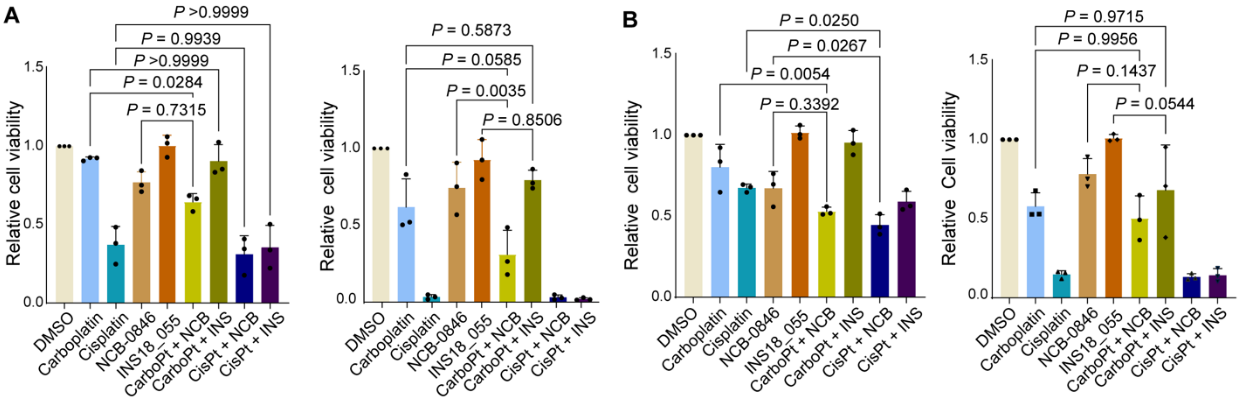


Hamidi et al., Supp. Figure 4

**Supplementary Figure S4. Effect of combination of TNIK inhibitors and platinum-based chemotherapy in LUSC cells.**

**A, B,** Cell viability (crystal violet assay) of SW900 (A) and NCI-H520 (B) cells treated with TNIK inhibitors (100nM NCB-0846 or 1 µM INS18_055), alone or in combination with carboplatin (6 µM) or cisplatin (5 µM), under two regimens: (left) co-treatment with TNIK inhibitors and platinum-based chemotherapy for 72 h, or (right) treatment with chemotherapy for 72 h followed by addition of the respective TNIK inhibitor for an additional 72 h. Data represent mean ± SD from *n* = 3 independent experiments; one-way ANOVA, Tukey's multiple comparisons post-test.

**Supplementary Tables**

**Supplementary Table 1. List of oligonucleotides used in this study (not described in material and methods).**

| | |
|---|---|
| *TNIK* qPCR | Forward: 5’ TCCACCAAAGGTGCCTCAAA 3’<br>Reverse: 5’ CCCAGAGCACTACCATTCCC 3’ |
| *MYC* qPCR | Forward: 5’ GGCTCCTGGCAAAAGGTCA 3’<br>Reverse: 5’ CTGCGTAGTTGTGCTGATGT 3’ |
| *RPL19* qPCR | Forward: 5’ TCGCCTCTAGTGTCCTCCG 3’<br>Reverse: 5’ GCGGGCCAAGGTGTTTTTC 3’ |
| *SNAI2* qPCR | Forward: 5’ CGAACTGGACACACATACAGTG 3’<br>Forward: 5’ CTGAGGATCTCTGGTTGTGGT 3’ |

**Supplementary Table 2. Clinical compound library with targets.**

| COMPOUND | REPORTED TARGET |
|---|---|
| Hydroxyurea | DNA replication |
| Tazemetostat | EZH2 |
| Pemetrexed | Purine and pyrimidine synthesis |
| Raloxifene (hydrochloride) | Estrogen receptor modulator |
| Mifepristone | Anti-progestogen & Anti-glucocorticoid |
| Fulvestrant | Anti-estrogen & selective estrogen receptor degrader |
| Enzalutamide | Androgen receptor antagonist |
| Rucaparib | Poly (ADP-Ribose) Polymerase-1 |
| 5-Fluorouracil | Thymidylate synthase |
| Gemcitabine | Nucleoside analog, DNA replication chain -terminator, ribonucleotide reductase |
| Erlotinib | Epidermal growth factor receptor |
| Saracatinib | Src and Bcr-Abl dual kinase inhibitor |
| Ipatasertib | Akt |
| Vorinostat | Histone deacetylase |
| Buparlisib | Pan-class I phosphoinositide 3-kinase (PI3K) |
| Everolimus | FKB12 |
| Topotecan(Hydrochloride) | Topoisomerase-I |
| Bortezomib | Proteasome |
| FRAX597 | Group I p21-activated kinases (PAKs) |
| Ganetespib | Heat shock protein 90 |
| Afatinib | Epidermal growth factor receptor, pan ERBB |
| Volasertib | Polo-like kinase |
| PF-573228 | Focal adhesion kinase |

| | |
|---|---|
| ART558 | DNA polymerase theta |
| Mirin | Mre11-Rad50-Nbs1 (MRN) complex |
| Ferrostatin-1 | Inhibitor of erastin-induced ferroptosis |
| Barasertib | Aurora kinase B |
| Infigratinib | Fibroblast growth factor receptors |
| Prexasertib | Checkpoint kinase 2 |
| Silmitasertib | Casein kinase II |
| Olaparib | Poly (ADP-Ribose) Polymerase |
| T0070907 | Peroxisome proliferator-activated receptor gamma (PPAR gamma) |
| Dabrafenib | BRAF V600E |
| Selumetinib | Mitogen-activated protein kinase kinase 1 and 2 |
| Y-27632 | Rho-associated protein kinase |
| IWR-1 | Tankyrase |
| Apoptozole | Heat shock protein 70 |
| Gefitinib | Epidermal growth factor receptor |
| Paclitaxel | Microtubules |
| SGI-1027 | DNA methyltransferase 1 |
| PFK-015 | 6-Phosphofructo-2-kinase (PFKFB3) |
| Nedisertib | DNA-dependent protein kinase |
| Lapatinib | Epidermal growth factor receptor, HER2 (ERBB2) |
| Dinaciclib | Cyclin dependent kinases (CDK 1, CDK2, CDK5 and CDK 9) |
| TAK-580 | Type II Raf kinases (including BRAF V600E, wild-type BRAF and wild-type CRAF |
| KU-55933 | Ataxia-telangiectasia mutated kinase (ATM) |
| GSK2606414 | Protein kinase RNA-like endoplasmic reticulum kinase (PERK) |
| Tipifarnib | Farnesyl protein transferase |
| Axitinib | Vascular endothelial growth factor receptors -1,-2,-3 (VEGFR) |
| Laduviglusib | Glycogen synthase kinase 3 beta (GSK3B) |
| Berzosertib | Ataxia telangiectasia and rad3-related (ATR) kinase |
| Ceritinib | Anaplastic lymphoma kinase (ALK) |
| Navitoclax | B-cell leukemia 2 (Bcl-2) family |
| Ro-3306 | Cyclin dependent kinase -1 |
| Empesertib | inhibitor of Monopolar spindle 1 kinase (Mps1) |
| SGX-523 | Hepatocyte growth factor receptor inhibitor |
| Alisertib | Aurora A kinase |
| Onvansertib | Polo-like kinase 1 |

| | |
|---|---|
| AZD1390 | Ataxia telangiectasia mutated (ATM) kinase |
| Camptothecin | Topoisomerase 1 |
| Tomivosertib | Mitogen-activated protein kinase (MAPK)-interacting serine/threonine kinase -1, -2 |
| FRAX1036 | p21-activated kinase 1 (PAK1) |
| Trametinib | Mitogen-activated protein kinase kinase 1 and 2, mutant BRAF |
| Lonafarnib | Farnesyl transferase |
| Niclosamide | Androgen receptor variant V7 |
| BML-277 | Checkpoint kinase 2 (Chk2) |
| Ceralasertib | Ataxia telangiectasia and rad3-related (ATR) kinase |
| ZN-c3 | Wee1 kinase |
| (+)-JQ-1 | Bromodomain-containing protein 4 (BRD4) |
| Adavosertib | Wee1 kinase |
| Adapalene | Retinoic acid (RAR) and retinoid X (RXRs) receptors |
| LB-100 | Protein phosphatase 2A (PP2A) |
| Palbociclib (monohydrochloride) | Cyclin dependent kinase -4 -6 |
| Copanlisib (dihydrochloride) | Phosphoinositide 3-kinase (PI3K) |
| Oxaliplatin | DNA cross linker |

### Supplementary Material and Methods

#### Cell lines

NCI-H520 (RRID: CVCL_1566) and SW900 cells (RRID: CVCL_1731) were obtained from ATCC; Lc-1sq (RRID: CVCL_3008) and LK2 (RRID: CVCL_1377) cells were obtained from the Japanese Collection of Research Bioresources (JCBR). All cell lines were verified by short tandem repeat profiling by IDEXX Bioanalytics. All cell lines were maintained at 5% CO2 at 37°C and cultured as recommended by the vendor and described [1]. In brief, NCI-H520 and SW900 were cultured in RPMI-1640 (Gibco, cat. 21870092) + 10% FBS (Tet-tested, heat-inactivated, Bio-Techne, cat. S10350H) + GlutaMAX (Gibco, cat. 35050061) + 1% penicillin–streptomycin (pen-strep; Gibco, cat. 15140122). Cells were tested for absence of Mycoplasma contamination by PCR every three months using the Mycoplasma Detection kit (Southern Biotechnology, cat. 103259-632).

#### Reagents

NCB-0846 (cat. S8392) and INS018-055 (ISM001-055; cat. E1944) were purchased from Selleck Chemicals (Houston, TX). NCB-0970, an inactive analog of NCB-0846, was synthesized in-house as described [2]. Doxycycline (cat. D3072) was purchased from Sigma-Aldrich (St. Louis, MO). Cisplatin (cat. HY-17394), carboplatin (cat. HY-17393), and tazemetostat (cat. HY-13803) were purchased from MedChemExpress (Monmouth Junction, NJ).

**siRNA-mediated transient knockdown**

SW900 or NCI-H520 cells (2 × $10^5$ cells/well in a 6-well plate) were reverse-transfected with corresponding siRNA oligonucleotides (50 nM) using Lipofectamine RNAiMAX (Invitrogen, cat., 13778150) according to the manufacturer's protocol. The medium was replaced 16 hours after transfection, and cells were analyzed for protein expression or subjected to functional assays (migration, invasion, senescence, EdU) at 72 or 96 hours after transfection. siRNA against MYC (SMART-pool, cat. L-003282-02) and non-targeting control siRNA #1 (cat. D-001910-04-20) were purchased from Dharmacon/Horizon Discovery (Lafayette, CO).

**Plasmid transfection**

The FUW-tetO-hMYC (Addgene #20723) plasmid was a gift from Rudolf Jaenisch; the FUW-tetO-MCS (Addgene #84008) was a gift from Stefano Piccolo. Plasmids were transfected into cells (2×105 cells/well in 6-well plates) using JetPRIME (Polyplus-Sartorius, cat. 114-15), and cells were analyzed 48 to 72 hours post-transfection.

**RNA extraction**

Genomic DNA was removed, and RNA was prepared using the RNeasy kit (Qiagen, cat. 74104) according to the manufacturer's protocol. RNA quantity and quality were determined using a NanoDrop ND-1000 spectrophotometer (NanoDrop Technologies, RRID: SCR_016517).

**RNA-seq analysis of TNIK-depleted LUSC cells**

Total RNA was extracted and purified using the RNeasy kit (Qiagen, cat. 74104) from SW900, NCI-H520, LK-2, Lc-1-sq, and HCC15 cells expressing a doxycycline-inducible control or TNIK-targeting shRNA (72 hours). RNA quality was assessed using a 2100

Bioanalyzer RNA 6000 Nano assay (Agilent, RRID:SCR_018043). RNA concentration was measured using a Qubit 2.0 Fluorometer (Life Technologies, RRID:SCR_020553). Illumina sequencing libraries were constructed using the NEBNext Ultra II Directional RNA Library Prep Kit for Illumina (NEB) and sequenced on Illumina NovaSeq 6000 (Illumina, RRID:SCR_016387) by pair-end sequencing with a read length of 2 × 150 bp by Novogene.

Sequence reads from RNA-seq experiments were aligned to the bowtie-indexed human HG38 genome with STAR [3]. The raw counts for each known gene from the RefSeq database were generated using htseq-count from the HTSeq package [4]. Differential expression between samples and across conditions was assessed for statistical significance using the R/Bioconductor package DESeq2 [5]. Genes with a false discovery rate (FDR) ≥ 0.05 and a fold-change ≥ 2 were considered significant.

The enriched canonical pathways and gene interaction networks of significant genes between shRNA control and shRNA TNIK#1 samples were generated using IPA (QIAGEN Inc.). The conditional hypergeometric method implemented in the GOstats package [6] was used to identify enriched Gene Ontology (GO) functional categories among genes of interest with a P-value cutoff of 0.001.

**Analysis of cell cycle distribution**

The Click-iT® Plus EdU Flow Cytometry Assay kit (Thermofisher, cat. C10420) and FxCycle™ Far Red Stain (Invitrogen, cat. F10348) were used to assess cell proliferation and cell cycle phase distribution. 1 × $10^5$ cells/well were seeded into 6-well plates. Following the indicated treatments, cells were incubated with EdU (10 µM) for 1 hour. Cells were then harvested, and EdU incorporation was detected according to the manufacturer’s instructions. After EdU detection, cells were incubated with 1 µL of FxCycle™ Far Red stain (30 min). Samples were analyzed in a BD FACSymphony A5 SE analyzer and data processed in FlowJo (v.10 or higher).

**FACS analysis of E-Cadherin, EpCAM, vimentin, and CD44 expression**

After the corresponding treatments, cells were fixed in fixation buffer (BD Cytofix, BD Biosciences cat. 554655, 30 min) and permeabilized with ice-cold methanol (90% v/v, 10 min). Subsequently, cells were incubated with the corresponding antibodies or isotype controls prepared in antibody dilution buffer (0.5% w/v BSA, 0.02% w/v $NaN_3$ in PBS). Antibodies against human CD324 (E-Cadherin) PE/Cyanine7 (cat. 324115), mouse/human CD44 [IM7] Pacific Blue™ (cat.103019), and anti-human CD326 (EpCAM) PE/Cyanine7 (Cat. 324221) were obtained from BioLegend (San Diego, CA) and used at the dilution recommended by the manufacturer. Antibody against human Vimentin Alexa Fluor® 488 (cat. 562338) was obtained from BD Biosciences (currently Waters Biosciences, Franklin Lakes, NJ). Samples were analyzed using a BD FACSymphony A5 SE analyzer, and data were processed in FlowJo (v.10 or higher).

**Cell Lysis and Immunoblots**

Whole-cell extracts were prepared by lysing the cells on ice in RIPA (Sigma-Aldrich, cat. R0278) lysis buffer supplemented with protease and phosphatase inhibitors (Halt Protease and Phosphatase Inhibitor Cocktail 100X, ThermoFisher, cat. 78446). Lysates were cleared (15,000 × g, 15 minutes, 4°C), after which protein concentration was determined with the Pierce BCA Protein Assay Kits, (Thermo Scientific, cat. 23227). Lysates containing equivalent amounts of protein were resolved by sodium dodecyl sulfate–polyacrylamide gel electrophoresis (SDS-PAGE) in 4% to 20% gradient gels (Bio-Rad, cat., 4561094), transferred into low-fluorescence polyvinylidene fluoride membranes, and membranes were blocked with Intercept Blocking Buffer (LICORbio, cat 927-60001).

Antibodies against TNIK (HL1751, cat. AB308194, 1:1000) and c-MYC (Y69, cat. ab32072, 1:1000, RRID:AB_731658) were obtained from Abcam (Waltham, MA). Antibodies against E-Cadherin (24E10, cat. 3195, 1:1000, RRID:AB_2291471), vimentin (D21H3, cat. 5741, 1:2000, RRID:AB_10695459), b-catenin (#D10A8, cat. 8480, 1:1000, RRID:AB_11127855) Phospho-p38 MAPK (Thr180/Tyr182) (D3F9, cat. 4511, 1:1000, RRID:AB_2139682), SOX9 (D8G8H, cat. 82630, 1:1000, RRID:AB_2665492), Histone H3 (D1H2, cat. 4499, 1:5000, RRID:AB_10544537), Slug (C19G7, cat. 9585, 1:1000, RRID:AB_2239535), GAPDH (14C10 cat. 2118, 1:2000, RRID:AB_561053), b.actin (8H10D10, cat. 3700, 1:5000, RRID:AB_2242334) were obtained from Cell Signaling Technology (Danvers, MA). Secondary antibodies, IRDye 800CW goat anti-rabbit IgG (cat. 926-32211, RRID:AB_621843) and IRDye 680RD goat anti-mouse (cat. 926-68070, RRID:AB_10956588) were obtained from LICORbio (Lincoln, NE). Blots were scanned using an ODYSSEY DLx imaging system (LICORbio).

**Cell fractionation assay**

SW900 or NCI-H520 cells (1 × $10^5$ cells/well in 6-well plate format) were seeded and treated with NCB-0846, NCB-0970, or INS018-055 for 72 hours. Cells were then harvested, and subcellular fractionation was performed using the Cell Fractionation Kit (Cell Signaling Technologies, cat. 9038) according to the manufacturer's instructions. Protein concentrations in cytosolic, nuclear, and membrane fractions were quantified and prepared for immunoblot analysis, as described above, alongside whole-cell extracts.

**Crystal violet cell viability assays**

Crystal violet cell viability assays were conducted and analyzed as described previously [1].

**C12FDG senescence staining assay**

Cells (5 × $10^4$ cells/well) were seeded in 6-well plates and treated with doxycycline (1 mg/ml; 96 hours) to induce shRNA expression. Cells were then incubated with C12FDG (5-Dodecanoylaminofluorescein Di-β-D-Galactopyranoside; Thermofisher, cat. D2893) according to the manufacturer's protocol. Senescent cells were identified by fluorescence-activated cell sorting (FACS) using a BD FACSymphony A5 SE Analyzer. Data were analyzed with FlowJo (Version 10 or higher, RRID:SCR_008520).

**Migration and invasion assays**

Corning Matrigel growth factor reduced basement membrane matrix (Corning, cat. 356234) was diluted in serum-free medium to a final concentration of 200 μg/mL. Next, 100 μL of the diluted Matrigel matrix was carefully added to the center of each Transwell® insert (8 μm PET membrane, Corning, cat. 3464) for invasion assays. Transwell inserts used for migration assays were not coated with Matrigel.

Cells (1.5 × $10^5$ cells/well) were seeded in 6-well plates and treated with the corresponding compound the following day. After 24 h (for TNIK inhibitor treatment) or 72 hours (for shRNA induction), cells were trypsinized and resuspended in serum-free medium. The cells were counted and diluted to a concentration of 5 × 105/mL in serum-free medium. Next, 150 µL of the cell suspension was seeded into the upper chamber of each Transwell insert. 800 µL of culture medium supplemented with 10% FBS, used as a chemoattractant, was added to the lower chambers. Serum-free medium was used as a basal control for migration or invasion. The cells were cultured in a humidified incubator at 37°C with 5% CO2 for 16 hours. Transwell inserts were washed twice with cold PBS, and the cells in the top chamber were gently removed using moistened cotton swabs. Migrating or invading cells were stained with crystal violet for 10 min and imaged under a BX50F4 Olympus microscope. Cells were quantified using ImageJ (National Institutes of Health, RRID: SCR_003070).

**Senescence-associated β-galactosidase staining**

Cells (5 × $10^4$ cells/well) were seeded in triplicate in 6-well plates and treated with TNIK inhibitors or doxycycline the following day for 72 or 96 hours. Senescence-associated β-galactosidase activity was assessed using the Senescence β-Galactosidase Staining Kit (Cell Signaling Technologies, cat 9860S) according to the manufacturer's protocol. Senescent cells were imaged with an Olympus BX50F4 microscope or an RVL2-K2 ECHO/Revolve microscope and quantified using ImageJ (National Institutes of Health, RRID:SCR_003070).

**Organoids and trypan-blue exclusion viability assay.**

A xenograft-derived LUSC organoid (XDO-4242) was established from patient-derived xenografts at the University Health Network [7]. The cultures were maintained in advanced DMEM/F-12 medium (Gibco, cat. 12634-010) supplemented with 2 mM GlutaMAX (Gibco, cat. 35050061), 10 mM HEPES (Gibco, cat. 15630-080), 100 U/ml Antibiotic-Antimycotic (Gibco, cat. 15240-062), 1X B-27 Supplement (Gibco, cat. 17504-044), 1.25 mM N-Acetyl-L-cysteine (Sigma-Aldrich, cat., A9165), 100 nM SAG (Enzo, ALX-270-426-M001), 50 ng/ml recombinant human EGF (Corning, cat. 354052), 40 ng/ml recombinant human FGF-7 (PeproTech, cat. 100-19), 100 ng/ml recombinant human FGF-4 (PeproTech, cat. 100-31), 100 ng/ml recombinant Human Noggin (PeproTech, cat. 120-10C), 0.5 µM A83-01 (Tocris, cat. 2939), 250 nM CHIR 99,021 (Tocris, cat. 4423), and 10 µM Y-27632 (Selleck Chemicals, cat. S1049) in growth factor-reduced Matrigel

(Corning, cat. 356231). Cells were dissociated into single cells using TrypLE Express (Gibco, cat. 12605-10) and were passaged every 6-10 days. When indicated, inhibitors or platinum-based chemotherapeutics were added 48 hours after seeding, and organoids were treated for 48 hours with the indicated compounds or combinations thereof. Organoids were collected, washed with 1X phosphate-buffered saline (PBS, Gibco, cat. 10010049) and dissociated into single cells using TrypLE Express. Cells were resuspended in cold PBS + 10% FBS and mixed 1:1 (v/v) with 0.4% trypan blue (Invitrogen, cat. T10282), and viable and dead cells quantified using a Countess 3 automated cell counter (Invitrogen, RRID:SCR_026963).

**Small-molecule libraries, screening, data processing, and analysis**

*Libraries*. The custom compound set, termed the FCCC Clinical compound collection. is comprised of seventy-five compounds selected for clinical targets (Custom set, MedChemExpress USA, Monmouth Junction, NJ). Compounds (**Supplementary Table 2**) were stored and diluted in DMSO.

*Screening*. Cells ($3.5 \times 10^3$ cells/well) were seeded in 96-well plates (Corning, cat. 07-200-89) using a Wellmate bulk reagent dispenser (Thermofisher, Waltham MA). Twenty-four hours after cell plating, cells were treated with DMSO (vehicle) or 250 nM NCB-0846. Library compounds were transferred via pin-tool transfer, and combination drugs were added using the Wellmate dispenser. Compounds were transferred from library source plates to cells in 96-well plates using a CyBio Well-Vario liquid handler (Analytik-Jena, Jena, Germany) equipped with a 100-nl pin tool (V and P Scientific, San Diego, CA) to achieve final concentrations of 200 nM or 1 mM. Seventy-two hours after drug addition, cell viability was assayed with CellTiter-Blue (Promega, Madison, WI, cat. G8081) using a Perkin Elmer Envision multilabel plate reader.

*Data processing and analysis*. Plate reader data were processed using the cellHTS2 package in R [8]. Plate-level normalization was performed by the median of negative control wells on each plate. Differential response to drug treatment relative to vehicle was assessed using limma [9]. Normalized, log2-transformed well-level intensities were fitted to a linear model with a design matrix encoding vehicle and drug-treatment groups, and a contrast (Drug -Vehicle) was tested. Moderated t-statistics were computed by empirical

Bayes shrinkage, and p-values were adjusted for multiple testing using the Benjamini-Hochberg method to control the false discovery rate (FDR). For each screen, wells were ranked by FDR-adjusted p-value and by drug/vehicle score ratio to prioritize hits. Compounds with FDR ≤ 5% and ratio change ≥ 0.15 were considered hits.
